# Hyperosmolality tolerance of freshwater largemouth bass (*Micropterus salmoides*) to brackish at early ontogenetic stages: Eggs, embryos and yolk-sac larvae

**DOI:** 10.1101/2023.12.16.572023

**Authors:** Du Luo, Dingtian Yang

**Affiliations:** South China Sea Institute of Oceanology, Chinese Academy of Sciences, Guangzhou 510301, China; Pearl River Fisheries Research Institute, Chinese Academy of Fishery Sciences, Guangzhou 510380, China; University of Chinese Academy of Sciences, Beijing 100049, China

**Keywords:** Osmoregulation, Osmotic homeostasis, Acute toxicity, Median lethal time, Hatching rate, Dose-response model

## Abstract

Salinity is recognized as a pivotal factor limiting the migration of freshwater fish to brackish environments. The largemouth bass (LMB, *Micropterus salmoides*), a globally translocated freshwater fish, exhibits estuarine distribution, yet its hyperosmoregulatory capacity during early ontogenetic stages remains inadequately understood. To investigate the impact of freshwater salinization, a series of experiments were conducted in the Pearl River Delta, China. The study aimed to elucidate the osmoregulatory abilities of LMB eggs and embryos, assess the salinity toxicity on hatching, and explore the acute effects of hyperosmolality on yolk-sac larvae. Our results revealed that freshwater-activated mature eggs and naturally fertilized oocytes maintained nearly identical osmotic homeostasis, with diameters of 1.38 ± 0.068 mm and 1.37 ± 0.054 mm, respectively. Furthermore, both exhibited peak water excretion at a salinity of 15.0 ppt. Remarkably, a reduction in water permeability was observed in hyperosmotic environments. Spontaneous hatching rates increased from 27.5 ± 14.4% in the 1.0 ppt group to 75.1 ± 12.0% in the 6.0 ppt group under fluctuating temperature conditions. Yolk-sac LMB larvae consistently reduced survival time from 12.5 d at 1.0 ppt to 50.7 ± 2.1 min at 35.0 ppt. Similarly, more developed larvae also experienced a decrease in survival time. Logistic regression models fitting lethal time with salinity indicated a sharp decrease between 10.0 ppt and 20.0 ppt. These findings offer practical insights for predicting distribution patterns and enhancing aquaculture technology for LMB. Moreover, they may contribute theoretically to the broader understanding of the osmoregulatory mechanisms of freshwater fish.

## Introduction

Largemouth bass (LMB, *Micropterus salmoides*) is one of the most widely distributed fish species in the world through human-mediated transmission. Their distribution range extends to every continent except Antarctica and Australia (Tidwell et al., 2019). LMB serves not only as one of the most popular sport fish but also as an important food fish with an annual production of 0.5 million tons (Hussein et al., 2020). Generally, it is considered a purely freshwater fish with strong adaptability to the environment. However, it was reported that LMB populations can also occur in oligohaline portions of estuaries (Devries et al., 2015). This piscivorous fish is commonly found in the oligohaline coast of estuaries in North America (Norris et al., 2010). On the one hand, the prevalence of salinization is extensive, which potentially endangers freshwater ecosystems (Cunillera-Montcusí et al., 2022). On the other hand, the adaptability of LMB to brackish water presents opportunities for both aquaculture expansion and natural distribution. Salinity, as one of the most important environmental parameters, has a key role in shaping aquatic biodiversity worldwide (Andrea et al., 2019). Salinity change is one of the evident mechanisms explaining the variation in species biomass or abundance. LMB is one of the fish species that responds positively and directly to salinity decreases (de Mutsert and Cowan, 2012). Presence in oligohaline sections of estuaries was a well-accepted scenario for LMB adaptation to increasing salinity (Norris et al., 2010). In artificial culture of juvenile LMB, saline water was found to be better than freshwater when growth and body contents were taken into consideration (Yi et al., 2021). As nonnative piscivores, it may threaten estuary-dependent species (McCormick, 2013). Therefore, understanding LMB salinity tolerance is extremely helpful for aquaculture production, ecological risk prediction and species management.

Fish inhabiting brackish environments must be capable of tolerating sudden and drastic fluctuations in salinity (Meador and Kelso, 1990). In the coastal regions of the Atlantic Ocean and Gulf of Mexico in the United States, LMB populations exhibit a wide occurrence in brackish waters (Meador and Kelso, 1989). A survey conducted on the Neuse River in North Carolina revealed the presence of LMB in areas with salinities ranging from 0 to 10.26 ‰ (Keup and Bayless, 1964). Although wild LMB tend to avoid higher salinities, they can endure salinities approaching 10.0 ppt in estuaries (Lowe et al., 2009). Native LMBs have been reported to inhabit estuarine waters with salinities up to 16 ppt (Peterson, 1988). As a species associated with freshwater environments, they may have developed increased tolerance to iron perturbation in elevated salinities. Smaller LMBs were found to be less tolerant of saltwater than their larger counterparts during the fingerling stage (Keup and Bayless, 1964). Nevertheless, fingerlings of 9.0 cm LMB can thrive when reared in water with a salinity of 10.0 ppt (Lu et al., 2022). As early as 1964, the effect of sea-water concentration on the reproduction and survival of LMB was examined (Tebo and McCoy, 1964). Reproduction in LMB significantly decreased when raised in concentrations exceeding 10% sea water. Fertilized eggs can hatch at 20% sea water but not at 30% (Heidinger, 1976). However, the detailed variations in hyperosmotic toxicity to LMBs across a broad salinity range, particularly during their early life stages, remain to be elucidated.

Teleost fishes are osmoregulators with ability of maintaining a constant osmolality of body fluids (Kültz, 2012a). Research indicates that salinities have the potential to impede or disrupt the normal development of fish eggs and larvae (Ban et al., 2022). The embryonic development process encompasses key stages, commencing with fertilization and progressing through zygote formation, cleavage, blastula formation, gastrulation, blastopore closure, germ layer differentiation, somitogenesis, and culminating in hatching. Notably, during the early stages of zygote development, exocytosis of cortical alveoli results in an increase in osmotic pressure, leading to the influx of water. Subsequently, a perivitelline space is formed between the zona pellucida and vitelline membrane, which serves as a protective barrier for the developing embryo. Adult fish maintain blood osmolality through the combined osmoregulatory functions of their branchial chambers, digestive system, urinary organs, and skin. In teleosts, both embryonic and post-embryonic stages exhibit osmolality within the range of 240–540 mOsm/kg. However, during the early life stages, the osmoregulatory capacity is primarily attributed to cutaneous ionocytes (Fridman, 2020). Teleost fish experience significant changes in egg envelope hardness during fertilization (Wang et al., 2021). The hydration of oocytes is influenced by changes in water salinity, which affect water and ion transport processes (Segret et al., 2022). Specifically for LMB, their eggs are adhesive with a prolonged embryo development process compared to free eggs (Rizzo and Bazzoli, 2020). Their osmoregulatory strategy in response to the acute-phase of habitat salinity remains unknown. The cellular stress response has evolved to enhance salinity tolerance in certain euryhaline fish species (Evans and Kültz, 2020). The permeability of water across animal cell plasma membranes is significantly greater than that of ions and other solutes (Chara et al., 2011). Although the change mechanism of cell volume has been studied in the intestines of *Fundulus heteroclitus* (Marshall et al., 2002) and *Anguilla anguilla* (Lionetto et al., 2005), the regulatory process remains unknown (Takei and Hwang, 2016). In the case of fish sperm, the adaptive capacity of its plasma membrane to drastic environmental changes is crucial for successful fertilization (Herrera et al., 2021). However, the response of osmolarity changes after exposure to hyperosmotic or hypoosmotic stress for fish eggs is seldom examined. Studies on the European eel have indicated that optimal egg diameter is achieved at salinities between 30 and 40 ppt, with a peak value observed at 35 ppt (Sørensen et al., 2016). In insects, an increase in osmotic pressure is required to trigger egg activation and volume expansion (York-Andersen et al., 2021). However, there is a lack of research investigating the direct effects of altered osmolarity on LMB eggs and fertilized zygotes cultured in saline water. Additionally, the specific impacts of salinity on LMB hatching and the potential lethal effects on yolk-sac larvae have not been elucidated. To clarify the salinity tolerance of LMBs at their early life stages, this study primarily aimed to determine the effects of freshwater salinization on the osmotic variation of eggs and zygotes. Secondly, we explored the influence of hatching naturally spawned embryos in brackish water, and thirdly, we examined the salinity tolerance of yolk-sac larvae.

## Materials and methods

### Osmolality variation of eggs and embryos exposed to saline media

Sexually mature LMB were procured from the Li Lang market in Guangzhou in April 2022. Mature eggs were carefully extracted from the abdomen and utilized in the experimental procedures. To create gradient salinities, a mixture of clean groundwater and sea salts was employed. Morphological changes, specifically variations in diameter, were utilized to assess the alterations in plasm osmolality of eggs when directly exposed to gradient saline solutions with salinities ranging from 1.0 ppt to 40.0 ppt. In each experiment, a quantity of 200 to 500 eggs from a single fish was introduced into a petri dish, previously filled with the respective gradient saline media, and exposed for one hour. Control experiments were conducted using eggs immersed in fresh groundwater. The room temperature was maintained within the range of 20 to 23 °C using an air conditioner. The morphology of these eggs was recorded using a microscope and digital camera. Subsequently, the length and width of individual eggs were quantified utilizing the sophisticated analysis software ImageJ (Rueden et al., 2017). Comparative analyses were conducted to assess morphological variations across distinct saline groups. The length and width of eggs from various saline groups were compared with each other to test if the difference was significant. Similarly, fertilized zygotes were collected from an artificial nest with natural spawning in aquaculture ponds. Then, these embryos (100 – 200 ind.) were incubated with fresh groundwater and gradient saline solutions with salinities of 1.0 ppt, 3.0 ppt, 5.0 ppt, 7.0 ppt, 9.0 ppt, 11.0 ppt, 13.0 ppt, 15.0 ppt, 20.0 ppt, 25.0 ppt, 30.0 ppt and 35.0 ppt. With the same methodology as previously described, the fertilized eggs cultured with gradient saline water for one hour and 18 h were observed, recorded, measured and analyzed. The experiments were conducted in triplicate.

### Hatching of fertilized eggs with artificial saline water

We conducted an experiment to further investigate the impact of salinization on the hatching process of LMB. Natural spawning eggs were collected from aquaculture ponds in Foshan within 6-15 h after fertilization in artificial nests. Graduated saline solutions, ranging from 1.0 ppt to 8.0 ppt, were prepared using sea salts and groundwater. Subsequently, 1500 ml of each solution was dispensed into plastic cups with a maximum capacity of 2000 ml. The experiments were replicated three times to ensure consistency and reliability of the results. Oxygen was supplied to each cup. The fertilized eggs were divided into saline media by cutting the egg-sticked windmill palm mats into small squires of 5×3 cm. To simulate the natural hatching process, we carried out the experiment on a farm to maintain the hatching water temperature changing with natural air temperature. The cultivating water temperature changed nearly with the field during the hatching process. The water temperature was recorded. Hatching was observed and recorded every four hours during the process. LMB hatching lasted for a relatively long time even under experimental conditions. We recorded spontaneous numbers of swimming larvae, surviving zygotes and obvious dead zygotes when most of the eggs hatched out.

The hatching rate (HRT) of per trial was calculated as follows:

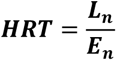

where L_n_ is the number of hatched larvae in each trial and E_n_ is the number of all the eggs used in each trial.

### Acute toxicity of yolk-sac LMB larvae exposed to gradient saline water

We used larval survival time as an indicator of hyperosmolality tolerance to acute toxicity. Yolk-sac LMB larvae were brought from fish farms in Foshan city to the laboratory and cultured using groundwater at a stable room temperature of approximately 20.0 °C. Series of saline solutions with concentrations of 1.0 ppt, 3.0 ppt, 5.0 ppt, 7.0 ppt, 9.0 ppt, 11.0 ppt, 13.0 ppt, 15.0 ppt, 20.0 ppt, 25.0 ppt, 30.0 ppt, 35.0 ppt and 40.0 ppt were separately divided into petri dishes with a diameter of 5.0 cm. Two-day-old larvae (total length = 0.5 - 0.6 cm) were slightly absorbed with plastic tubes and released directly into each petri dish. To clearly observe the lethal process and record the dead larvae, only six to eight ind. larvae were released into each dish. When there was no movement behavior with slight shaking and soft touching, the individuals were recorded as dead. At the same time, the survival time was also recorded. Approximately 86.0 h later, the larvae that were cultured with the same temperature and air supply conditions in groundwater were put into 7.0 ppt to 40.0 ppt saline solutions with the same methods to investigate the lethal effect and to make a comparison with the previously exposed two d larvae. The middle lethal time and lethal time of each trial were recorded. The experiments were conducted in triplicate.

### Statistical analysis

For the treatments without direct records, we calculated the median lethal time through online tools based on probit analysis according to Vinay Kumar et al. (http://14.139.232.166/probit/probit-analysis.html) (Kumar et al., 2020). Before the analysis and visualization of the size variation of eggs and embryos, we standardized all the mean values by using z score standardization through the scale function with the dplyr package in R. The standardized data were used to perform linear and nonlinear regression to explore variations with increasing water salinity. In the hatching experiment, normality and lognormality were tested for all data from different salinity groups before ANOVA for significant differences. All the data are given as the mean ± standard deviation. Hatching rate and survival time among various salinities were compared by ANOVA (*p* = 0.05) and multiple range test (*p* = 0.05). The hatching success of fertilized eggs was analyzed with the aov() function and Tukey’s HSD test for multiple comparisons of means. The regression relationship between salinity and lethal time was determined using the drc package in R software. In scattering plots, the regressed line was smoothed using loess in R. Comparison of zygote diameter variation and larvae survival time between groups at two different times was performed using paired t tests. All statistical analyses and visualizations were performed using R 4.1.1 (R Core Team, 2022)

## Results

### Osmolality variation of prespawn LMB eggs exposed to saline water

In order to investigate alterations in osmolality, we assessed the size differences in both mature and fertilized LMB eggs, using these variations as indicative measures. The analysis of data derived from 14 fish revealed that the average diameter of freshly matured fish eggs was 1.24 ± 0.093 mm, with a range spanning from 0.53 mm to 1.73 mm. A total of 8,394 mature eggs from nine fish were measured to test the size variation of eggs cultured in a series of saline waters with salinities ranging from 0 to 40.0 ppt. We tested an average of 68 eggs for each saline level from every fish. Among all the treated fish groups, the largest eggs appeared in one freshwater group with a diameter of 1.47 ± 0.15 mm. The smallest eggs were 1.11 ± 0.07 mm in another group with a salinity of 9.0 ppt. When all the eggs with diameters > 1.0 mm from nine fish were taken into consideration, the average diameter of LMB eggs was the largest at 1.38 ± 0.068 mm in freshwater, and the smallest eggs were 1.23 ± 0.066 mm in 15.0 ppt brackish water (Fig. 1A). This salinity of 15.0 ppt was also the predicted concentration where the eggs shrined most with water excretion. As if measured by percent, the egg diameter increased by 8.35% in freshwater and decreased by 3.43% in 15.0 ppt saline water.

**Fig. 1.**
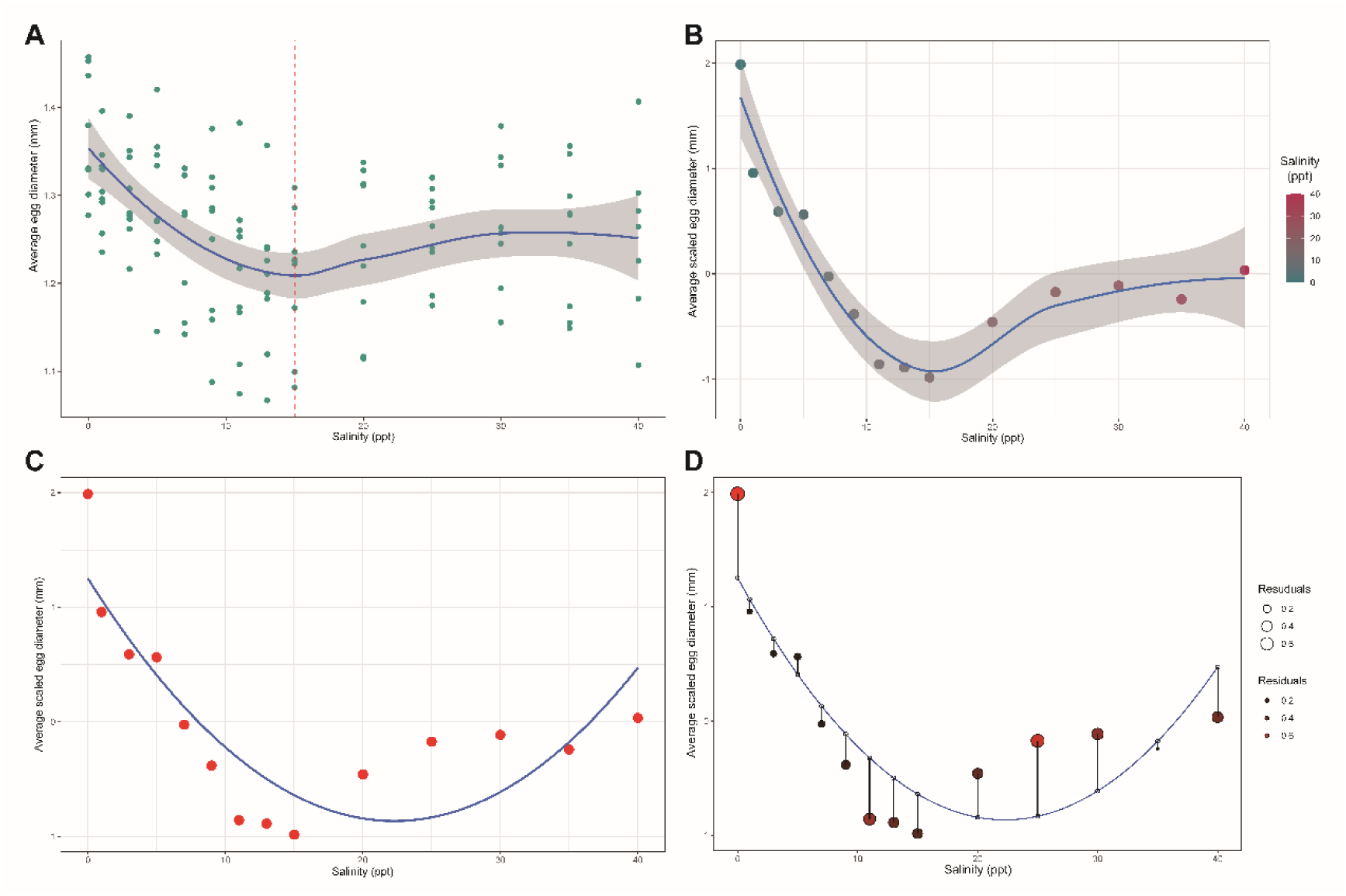
Size variation of the mature eggs maintained in a series of saline water with concentrations ranging from 0 to 40.0 ppt (**A**). The regressed smooth line with a 95% confidence interval for a mean represented the change in the average egg diameters with increasing salinity. The vertical dotted line shows the salinity of 15.0 ppt, which was the predicted concentration where the largest size variation disappeared. Size variation of the mature eggs indicated by a smooth line regressed with the loess method based on average scaled diameter data (**B**). The level of confidence is 95%. Relationship between salinity concentration and scaled egg diameters regressed with quadratic polynomials (**C**). Residuals between scaled test egg diameters and corresponding predicted data (**D**). The length of the bar represents the difference between the actual diameter and the predicted line.

As the size of fresh eggs from different fish changed greatly among themselves, we scaled the morphological measurements of 0 - 40.0 ppt saline water-treated eggs from each fish. The predicted size variation line based on average variation indicated that the egg diameters declined first when the salinity concentration varied from 0 to 15.0 ppt and then rose back with increasing salinity (Fig. 1B). Therefore, we regressed the nonlinear variation using scaled average data with a potential quadratic polynomial formula (Figure 1C). The regressed formula was y = 0.0043x^2^ - 0.19x + 1.25. where x is the salinity concentration and y is the scaled egg diameter. It fitted relatively well with scaled test data with a correlation index of multiple R^2^ = 0.73. At salinities of 0 - 5.0 ppt, the egg diameter increased over 2.0%. However, at salinities of 9.0 - 20.0 ppt, the egg diameter decreased by over 1.0%. There was no obvious variation in other tested salinity levels at 7.0 ppt and 25.0 - 40.0 ppt.

We analyzed the difference in residuals and residuals between the predicted data and test data (Fig. 1D). The results suggested that the model biased the variation in egg diameters mainly at salinity concentrations of 0, 11.0 and 25.0 ppt. When the diameters of eggs immersed in various saline water were regressed with the segmental salinity concentration, the selected higher correlation index (R^2^) was 0.91 and 0.95, respectively, with line regression and dimensional multiple regression between 0 - 15.0 ppt. The change in egg size was highly correlated with water salinity when regressed with dimensional multiple models in salinity ranging from 15.0 ppt to 40.0 ppt (R^2^ = 0.91) and in salinity ranging from 15.0 ppt to 35.0 ppt (R^2^ = 0.99).

### Osmotic variation of fertilized eggs cultivated with saline water

The diameter of fertilized eggs was measured approximately 10 h post-fertilization, ranging from 1.27 mm to 1.48 mm, with an average of 1.37 ± 0.054 mm (n = 969). Subsequent to an 18-hour treatment, there was a substantial reduction in their diameters at salinity concentrations of 7.0 ppt, 9.0 ppt, 11.0 ppt, 13.0 ppt, and 15.0 ppt, in comparison to measurements taken after one hour of treatment (p < 0.001) (Fig. 2A). Additionally, notable variations were observed in the salinity groups of 25.0 ppt (p < 0.01) and 35.0 ppt (p < 0.05). Among these variations, the 15.0 ppt group was the greatest, followed by the 13.0 ppt group. The zygote diameter decreased 11.2% and 9.4% in the 13.0 ppt and 15.0 ppt groups, respectively. When scaled size variation of saline-treating for one hour and 18 h to starting, and one hour to 18 h were recorded, we found that there were slight changes in one-hour-treatment. However, 18 h of treatment greatly changed zygote size at the middle concentration levels, and all the salt-treated zygotes were shortened (Fig. 2B). Both one-hour treatment and 18-hour treatment showed fluctuations at different salinity levels. Regressed diameter variations showed that the size decreased greatly with increased salinity from 1.0 to 13.0 ppt after 18 h of culture (Fig. 2C). Saline water of 13.0 ppt and 15.0 ppt was the most capable of decreasing diameters. Linear regression indicated that the diameter decreased stably with increasing salinity from 0 to 15.0 ppt (R^2^ = 0.91). The size change was small when the salinity concentration was higher than 20.0 ppt. However, the variations in zygote diameters after treatment for one hour were not highly linearly correlated at all salinity concentrations or at lower salinity concentrations. The diameter decline was observed only at lower salinities of less than 9.0 ppt.

**Fig. 2.**
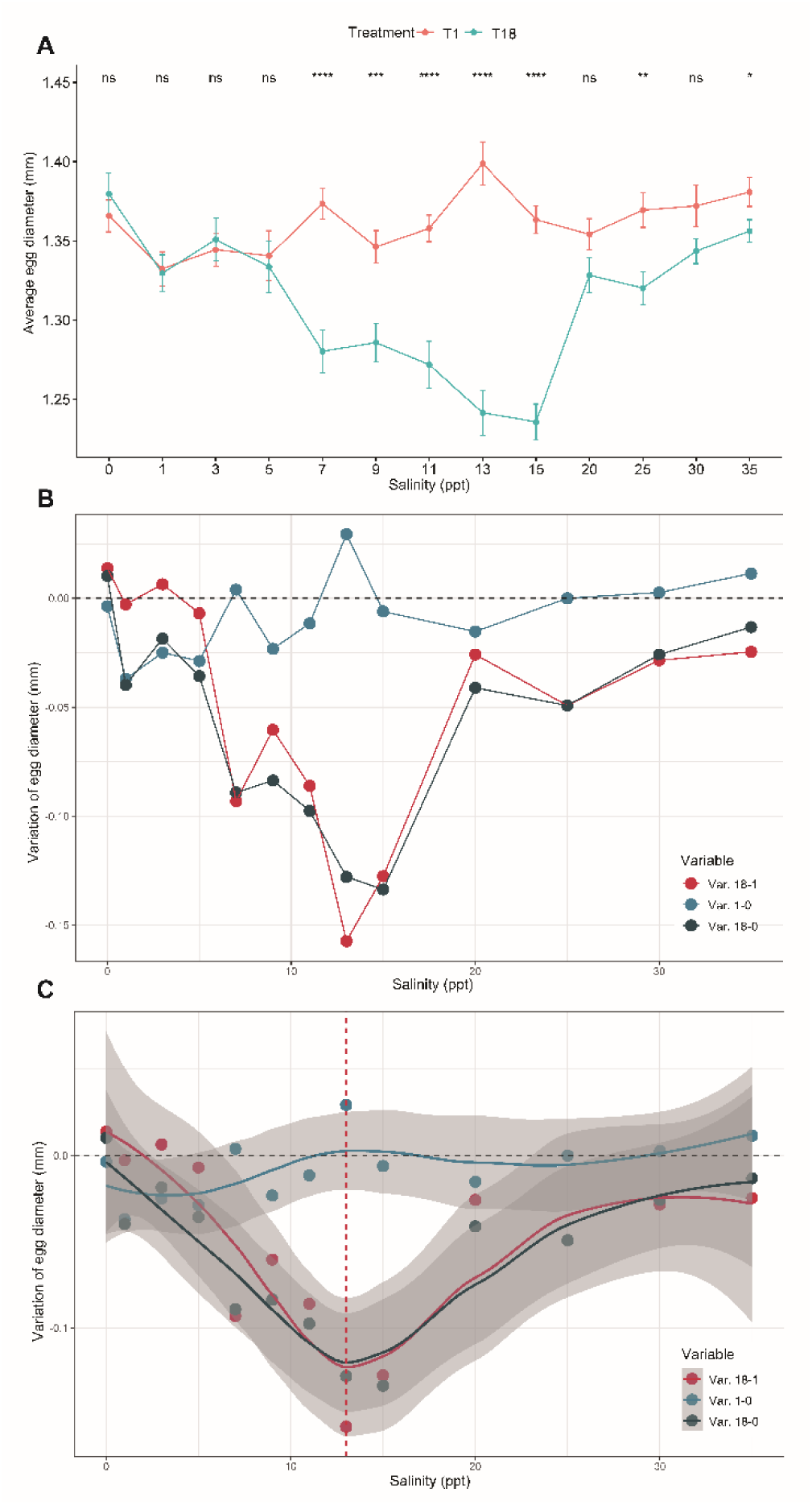
Comparisons of zygote diameter maintained with various saline water for one hour (T1) and 18 h (T18) for each concentration varying from 1.0 to 35.0 ppt (**A**). Freshwater as a salinity zero group was used as a control. Scaled size variation of zygotes cultured with 13 levels of saline water(**B**). Size variation was calculated between the first hour to the beginning (Var. 1-0), 18th hour to the beginning (Var. 18-0) and 18th hour to the first hour (Var. 18-1). The dotted line shows no diameter variation. Regressed diameter variation of zygotes (**C**). Size variation was calculated between the first hour to the beginning (Var. 1-0), 18th hour to the beginning (Var. 18-0) and 18th hour to the first hour (Var. 18-1). The dotted line shows the salinity concentration of 13.0 ppt.

### Hatching of fertilized eggs in saline water at fluctuating lower temperatures

The hatching process was conducted under semi-natural air temperature conditions. Hourly air temperature data were obtained by referencing Guangzhou as the target city and retrieving historical records online from https://www.timeanddate.com/ (accessed on 2022-10-6) for detailed temperature information. Then, we investigated the variation in environmental temperature during the hatching process. The air temperature experienced an M-type variation of two increases and two decreases during the experiment with a period ranging from Mar. 30^th^ to Apr. 2^nd^ in 2022 (Fig. 3). In the first half of the experiment, the air temperature increased from 18.0 °C to 30.0 °C; however, it decreased to 12.0 °C at the near end of the experiment. The hatching water temperature changed with the air temperature. We recorded the highest water temperature of 28.6 °C and the lowest water temperature of only 14.6 °C. The estimated average temperature was 19.8 °C.

**Fig. 3.**
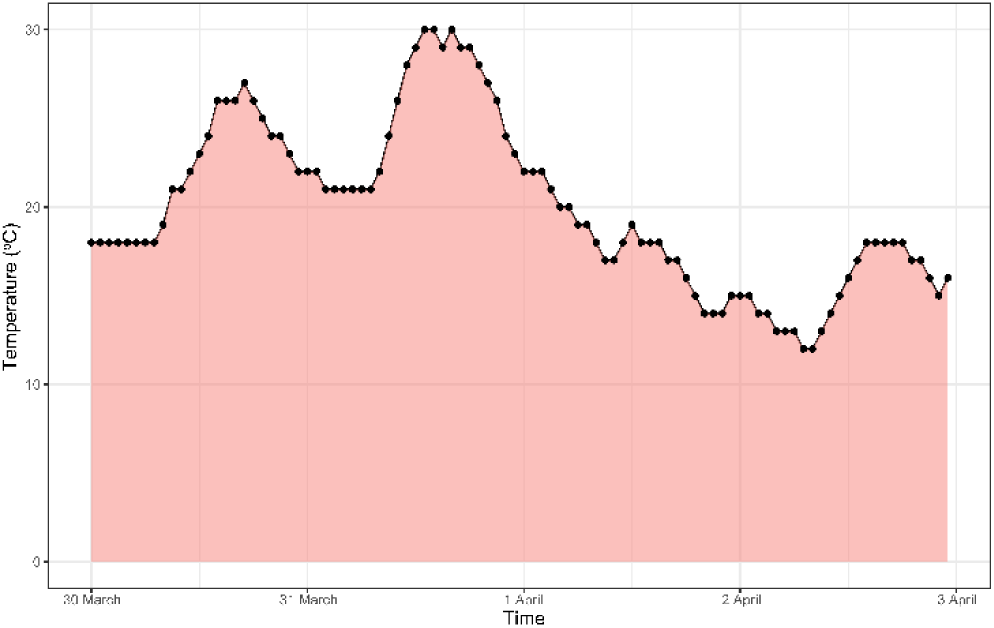
Variation in air temperature during the hatching process. The temperature was given in degrees centigrade (°C) hour by hour from 30 March to 3 April in 2022.

The Shapiro‒Wilk test was used to test the normal distribution of spontaneous hatching time, survival rate and lethal rate for each group with salinity ranging from freshwater to 8.0 ppt. The hatching experiment started at 19:00 on 30 March and ended at 21:00 on 2 April. We examined the spontaneous hatching rate (HRT), survival rate of embryos, and lethal rate of the zygotes after 75.0 h of hatching and approximately 84 h after spawning. The spontaneous hatching rate was 63.1 ± 4.9% in freshwater. In saline waters, the hatching rate stably increased with salinity from 1.0 ppt to 6.0 ppt (Fig. 4A). HRT in Groups 1.0 ppt (27.5 ± 14.4%) and 6.0 ppt (75.1 ± 12.0%) was the lowest and highest, respectively. HRT in the 7.0 ppt group (71.9 ± 14.6%) was close to that in the 6.0 ppt group, while it declined dramatically in the 8.0 ppt group (39.2 ± 10.9%). When groups of salinity = 2.0 ppt and 5.0 ppt, without normal distribution, were excluded from Tukey’s multiple comparisons test, we found that the spontaneous hatching rate of embryos from Group 1.0 ppt was significantly different from Group 6.0 ppt (*p* = 0.0075) and 7.0 ppt (*p* = 0.013).

**Fig. 4.**
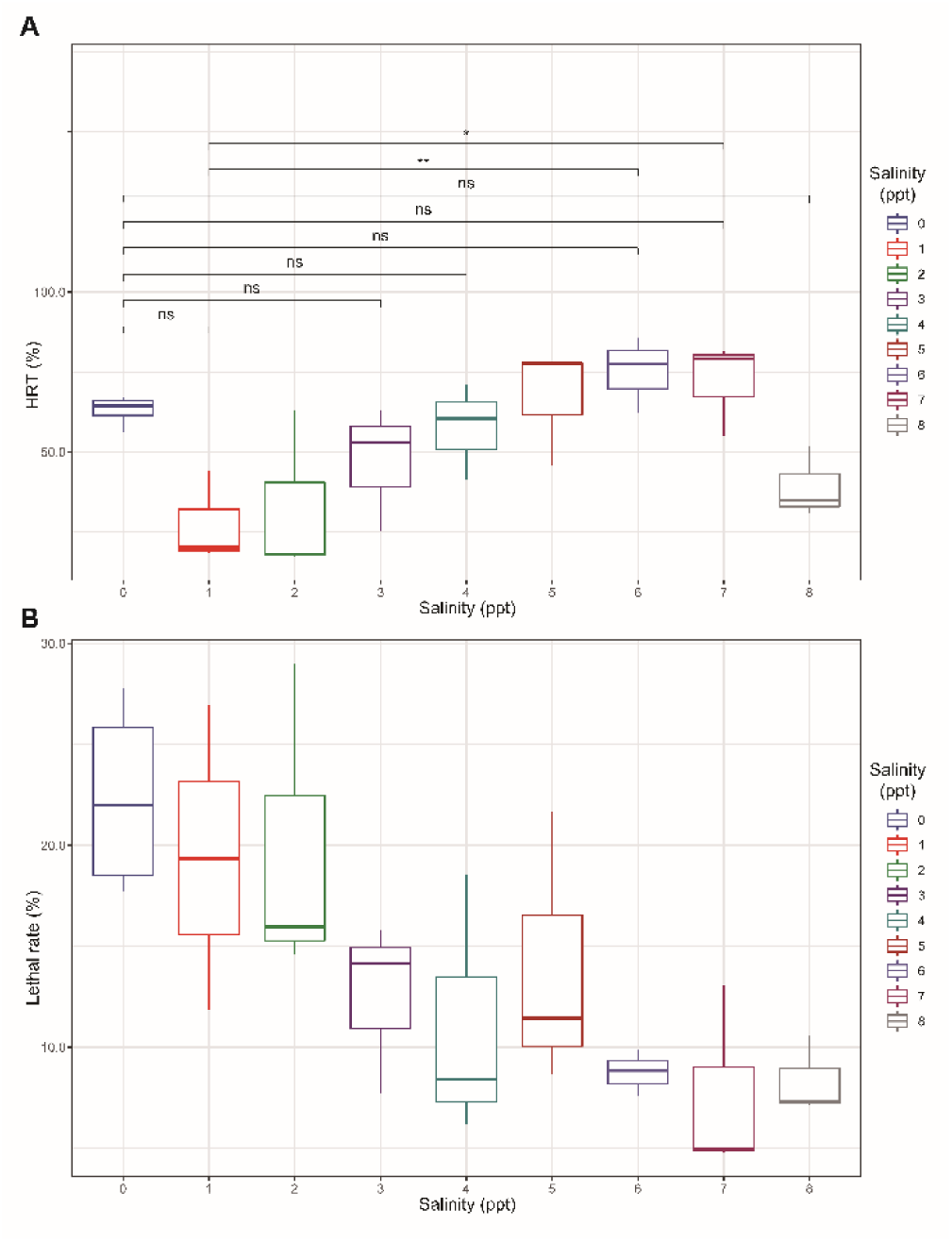
The spontaneous hatching rate (HRT) of the largemouth bass fertilized eggs hatched with various saline water with concentrations ranging from 0.0 ppt to 8.0 ppt (**A**). The box shows the 25th percentile, the median and the 75th percentile of HRT in each group. The length of the line represents the standard deviation. The symbol ns indicates no significant differences. * and ** indicate the difference between two groups. Lethal rate of the largemouth bass fertilized eggs hatched with various saline water with concentrations ranging from 0.0 ppt to 8.0 ppt (**B**). The box shows the 25th percentile, the median and the 75th percentile of the lethal rate in each group. The length of the line represents the standard deviation. We recorded the number of obviously white-colored dead eggs and calculated the percentage of these lethal zygotes. The lethal rate of 22.4 ± 4.9% was highest when hatching in freshwater (Fig. 4B). However, it decreased from 19.4 ± 7.6% in the 1.0 ppt group to 7.6 ± 4.7% in the 7.0 ppt group. Then, the lethal rate increased in the 8.0 ppt group. Taking all levels to make a line regression analysis, we found that the lethal rate constantly declined from 0.0 ppt to 8.0 ppt (R^2^ = 86.1).

### Acute toxicity of yolk-sac LMB larvae exposed to gradient saline water

Yolk-sac LMB larvae varied their tolerance to different concentrations of salinity. One group of larvae lived the longest time of 17960.0 min, approximately 12.5 d, in 1.0 ppt solution. It was even longer than the longest group with 17,741.0 min, approximately 12.3 d, in freshwater during the experiment. However, saline water with a salinity of 35.0 ppt led all the larvae to die in less than 50.7 ± 2.1 min. We smoothed the line of the mean full lethal time for each saline level with the loess method in R (Fig. 5A). When the newly hatched larvae were released into saline water with salinity contents ranging from 1.0 ppt to 35.0 ppt, their survival time decreased continuously with varying changing rates. A total of 52 dose‒response-related models in the package drc in R were used to fit the change in survival time with gradient saline contents. The dose-response model of BC.4 was relatively the most fit to the change in larvae survival time with *p* = 0.021 (Fig. 5B). Both the smooth line and the regression line showed that there was an extremely significant decrease between salinity concentrations of 10.0 ppt and 20.0 ppt. In particular, the average survival time decreased from 9,362.0 ± 1049.6 min at 11.0 ppt to 2,386.3 ± 86.4 min at 15.0 ppt, and the mean survival time was only 371.0 ± 106.5 min at 20.0 ppt. The predicted survival time was 1220.3 min at a salinity of 16.0 ppt.

**Fig. 5.**
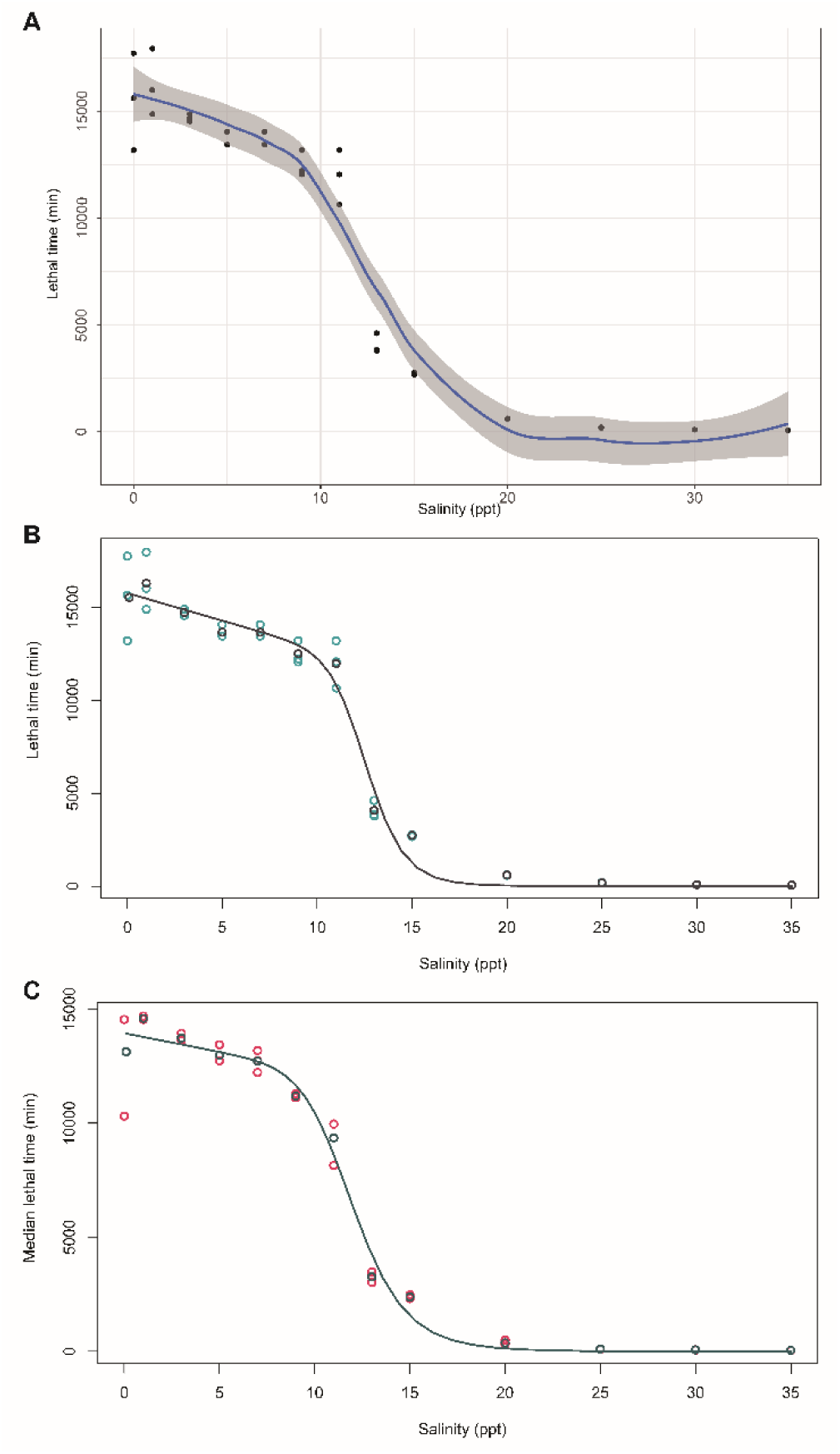
Smooth of the mean survival time of the yok-sac larvae treated accurately with different saline water with salt concentrations ranging from 0 ppt to 35.0 ppt (**A**). The confidence interval was 95%. The nonlinear regression line was depicted to imply the decreasing pattern of survival time (**B**) and the decreasing pattern of mean median lethal time (**C**) for various saline levels. The two kinds of points were the individuals and average experimental test under each saline level, respectively.

The median lethal time of 14595.0 ± 84.9 min in the 1.0 ppt group was also higher than that in the other groups. However, a longer median lethal time was not absolutely consistent with a longer full lethal time. The difference between median lethal time and lethal time for each concentration varied greatly from 2386.9 min in groups of fresh water and 1688.7 min in groups of 1.0 ppt to 1695.0 min and 2603.0 min in other saline groups of 9.0 ppt and 11.0 ppt, respectively. Although the gap between the median lethal time and lethal time was larger at lower saline concentrations, the variation rate appeared at higher concentrations of 25.0 ppt and 20.0 ppt. The model of BC.4 also fit well to the mean of the median lethal time. There was also an apparent decrease in the median lethal time in salinity between 10.0 ppt and 20.0 ppt (Fig. 5C).

At an average of 86.0 h later, the larvae from the same parent cultured in freshwater changed their salinity tolerance. The larval survival time declined at all the examined salinity levels from 7.0 ppt to 35.0 ppt (Fig. 6). The median lethal time and lethal time at a salinity of 40.0 ppt were only 12.3 ± 1.2 min and 15.8 ± 0.4 min, respectively. Comparing means of T tests showed that the larvae’s survival time significantly (*p* < 0.05) decreased in salinity concentrations from 7.0 ppt to 15.0 ppt, except groups of 9.0 ppt. When salinity increased from 20.0 ppt to 35.0 ppt, the variations in lethal time became extremely significant (*p* < 0.01).

**Fig. 6.**
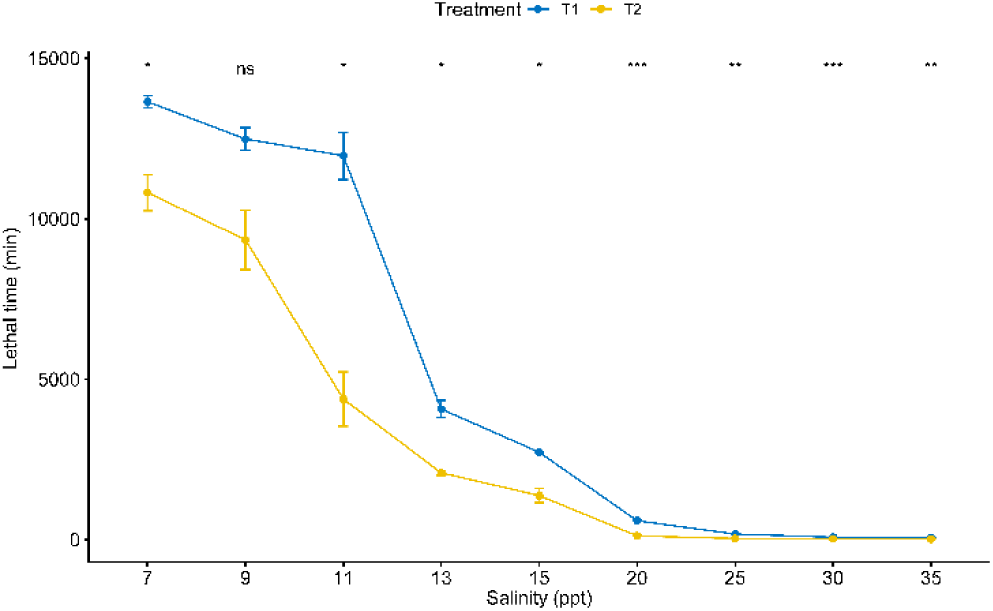
Mean survival time of the largemouth bass larvae maintained at various salinity levels. T tests were applied to detect the potential difference in the mean lethal time between the first treatment (44 h after hatching) and the second treatment (130 h after hatching) at each salinity level. The labels *, **, ***, and ns represent different significance levels of *p* < 0.05, *p* < 0.01, *p* < 0.001 and *p*>0.05, respectively.

## Discussion

Salinity emerges as a pivotal barrier hampering the migration of freshwater fish into brackish environments (Peterson and Meador, 1994), exerting significant control over the activity and distribution of aquatic species. The ability of euryhaline teleosts to regulate osmotic balance enables them to adapt physiologically to fluctuating environmental salinity (Kültz, 2015). The successful establishment of introduced fish species hinges on the comprehensive recruitment of offspring and subsequent growth, laying the groundwork for a self-sustaining population (Katelyn and Jeffrey, 2021). Furthermore, Investigating the adaptability of fish to varying osmotic pressures becomes crucial for the development of saline-alkali water fisheries. In this study, we probed the osmolality variation of LMB eggs cultivated in saline water across diverse salinity levels, both pre- and post-fertilization. Our investigation extended to assessing the tolerance of fertilized eggs to hatching in saltwater and exploring potential adverse effects of elevated salinity on yolk-sac larvae. The collective exploration sought to unravel the adaptive features and potential mechanisms governing early ontogenic stages of LMB.

Fish eggs varied in characteristics corresponding to reproductive strategies. In comparison to long-migratory reproductive species, LMB eggs may be less numerous and possess a degree of adhesiveness (Zelazowska and Halajian, 2019). Furthermore, LMB eggs display diversity in diameter and feature small perivitelline spaces. Adhesive eggs have surface specialization, and their size is associated with the structure of the zona radiata. It is still uncertain whether the distinct area of the plasma membrane at the base of the micropyle has osmotic adjusting capability. For fresh water fish, their fertility was limited to a few seconds after the ovulation of the oocytes were released into water. Previous studies have reported that the diameter of mature LMB eggs ranges from 0.75 to 1.5 mm (Tidwell et al., 2019). Our study found that the diameter of nonactivated mature eggs varies from 0.53 mm to 1.73 mm, which is larger than the previously reported range. However, there have been no reports about the fertilizing or hatching ability of small-sized LMB eggs until now, and most of the eggs were larger than 1.0 mm. Therefore, we excluded small eggs < 1.0 mm from the present investigation to examine volume variation.

The mature fish oocyte, upon spawning, comprises a substantial yolk mass enveloped by a thin peripheral layer known as the cortical lamina. Externally attached to the plasma membrane is the chorion, which consists of nine strata (Dumont and Brummet, 1980). Post-fertilization, the chorion undergoes structural changes, hardening to provide protection to the developing embryo. The cortical reaction, an essential step in egg activation, occurs in all teleost eggs, regardless of fertilization status. Throughout incubation, the diameter of the eggs exhibits only slight variations (Coward et al., 2002). In the context of our hatching experiment, the fertilized LMB eggs displayed an average diameter ranging from 1.27 mm to 1.48 mm. This observation suggests that the freshwater-induced egg activation during spawning significantly strengthens and enlarges the egg envelope. Further exploration, including comparisons between fertilized and unfertilized mature eggs, could enhance our understanding of the egg envelope’s role in responding to hyperosmotic stress.

When comparing pre-ovulating mature eggs to fertilized zygotes, it was observed that a salinity of 15.0 ppt predominantly led to a reduction in the diameter of both types of eggs. In brackish water, spanning a salinity range from 0 to 15.0 ppt, both pre-ovulating mature eggs and fertilized zygotes exhibited a linear decrease in size (R2 = 0.91). Furthermore, the average diameter of the freshwater-activated eggs was nearly the same (1.37 mm), regardless of whether the mature eggs were directly acquired from the fish abdomen or fertilized eggs from natural spawning. This indicated that the osmoregulatory ability would not be determined by the action of fertilization or the structure of the micropyle. In contrast to eggs, the transfer of eels directly from seawater to freshwater resulted in a significant reduction in plasma osmolarity in their somatic cells, particularly in the epithelial cells, with a maximum decrease of 18.6% (Lionetto et al., 2005). However, differences in osmoregulation were evident between the two types of eggs when incubated in saline water with a salinity of 20.0 ppt. Under this salinity level, mature LMB eggs experienced a reduction in diameter, while the size of fertilized zygotes remained relatively stable. No significant variation in egg size was observed when the incubation salinity exceeded 20.0 ppt. This scenario suggests that osmolality adaptation is an active process orchestrated by the egg envelope.

For euryhaline fish, the adaptation strategy to salinity may undergo a shift when the environmental salinity surpasses their upper tolerance limit (Kültz, 2015). Consistent with this study, previous research indicates that such shifts in osmoregulatory strategy are evident under both extremely high and extremely low salinity conditions. Under extremely hyperhaline conditions, water retention takes precedence over paracellular secretion of Na^+^, and the secretion of Na^+^ relies on leaky tight junctions (Kültz, 2015). In adhered cells, the measured osmotic pressure is demonstrated to be dependent on the number of proteins and ionic content (Adar and Safran, 2020). Evidence from the turbot *Scophthalmus maximus*, a species with multiple adaptations to various salinities, suggests that osmotic regulation is associated with various genes (Cui et al., 2020). However, pinpointing osmosensors in fish remains challenging due to the abundance of proteins that meet the functional criteria (Kültz, 2012b). Consequently, the mechanism underlying their osmotic regulation remains unknown (Wang and Kültz, 2017).

The embryonic development process may be affected by physical and chemical environmental characteristics of water bodies. In the development of zygote stages, exocytosis of the cortical alveoli increases the osmotic pressure and leads to water influx. Then, a perivitelline space between the zona pellucida and vitelline membrane was created, which functioned as embryo protection (Rizzo and Bazzoli, 2020). The present research found that it took nearly four days to hatch the eggs at an average temperature of 19.8 °C, ranging from 14.6 °C to 28.6 °C. It was close to the natural hatching time of three to four days (Tidwell et al., 2019). Similarly, in Lake Powell, the mean daily water temperature at nests increased from 14.4 °C to 23.9 °C during the spawning period in freshwater. All nests required four or five days to hatch and maintain the survival of embryos to hatching at 80.4% - 92.2% (Miller, 1971). Maintaining a consistent egg incubation temperature has been reported to enhance hatching rates and decrease dependence on chemical treatments (Matthews et al., 2012). However, the combined effects of temperature and salinity on hatching rates remain unknown.

The spontaneous hatching rate in freshwater (63.1 ± 4.9%) and the highest hatching rate in saline water (75.1 ± 12.0%) were both observed to be relatively lower than the normal fertilization rates reported in aquaculture reproduction (Tidwell et al., 2019). Lower temperatures were found to extend the hatching period of fertilized zygotes, and the presence of proteolytically active fractions capable of digesting the zona radiata interna may contribute to envelope digestion swelling (Schoots et al., 1983). This implies that incubating LMB eggs at lower temperatures and higher osmolality could adversely impact zygote development. Furthermore, the lower hatching rate and higher egg lethal rate at a salinity level of 8.0 ppt also suggest a potentially hazardous effect of osmolality on LMB oocytes. This result was consistent with a previous report that there were no hatched fertilized LMB eggs in diluted marine water with a salinity of 10.61 ppt (Tebo and McCoy, 1964). Although, lower temperature may decrease the eggs’ enzymatic activity and trigger additional risk from incubating water. Water salination could inhibit the growth of microorganisms hazardous to fish eggs (Tidwell et al., 2019). This was consistent with the findings that the zygote mortality rate stably declined with increasing salinity from 0 to 7.0 ppt. A previous report also mentioned that 4.0 and 6.0 ppt salt treatments were effective for controlling water mold to improve swim-up rates during the incubation of LMB eggs (Tidwell et al., 2019). Salination of rearing water would suppress the hazardous effect of microorganisms from the environment.

Osmoregulation serves as the fundamental mechanism enabling fish to adapt to alterations in environmental salinity. Our study demonstrated that LMB yolk-sac larvae could endure for over 12 d in lower salinity water. However, their survival in saline water with a salinity of 20.0 ppt was limited to approximately 6 h. The correlation between survival time and salinity level revealed a notable decline, particularly between 10.0 ppt and 20.0 ppt. Previous research suggests that freshwater fish typically tolerate and survive salinities less than 9‰ (Peterson and Meador, 1994). Despite being freshwater inhabitants, LMBs exhibited relatively robust adaptability to brackish water in our investigation. Yet, further exploration is crucial to understand the behavioral and physiological responses of LMBs within this salinity range. The determination of lethal effects is grounded in empirical connections derived from quantitative exposure-response data, while the intermediary processes remain inadequately understood (Ankley et al., 2010). Teleosts are likely to display a higher survival rate when exposed to acidic or alkaline waters in comparison to soft waters (Baldisserotto, 2019). Additionally, the impact of iron in the environment on the osmoregulation of LMB requires further exploration.

In its entirety, the complete lethal time and median lethal time of LMB larvae in response to varying salinity levels conformed well to the logistic model BC.4. However, the observed individual variability in survival time, as well as the disparities between full lethal time and median lethal time at each salinity level, suggested a degree of individual variation in adapting to higher osmolality. Our results suggest a significant decrease in larval survival time within the salinity range of 7.0 ppt to 15.0 ppt, indicating a potential inhibitory effect of salinity on LMB larval development. Moreover, a notable decline in survival time is observed at salinity levels exceeding 20.0 ppt, suggesting an intensified toxic effect of high osmolality, particularly for more developed individuals. However, whether this difference is attributed to metabolic factors or physical resistance remains unclear.

LMB may exhibit varying adaptability across different ontogenetic stages. A previous study reported that sac fry hatched in saline water displayed poor development and had a survival duration of 11.6 d and 9.8 d at salinity levels of 5.32 ppt and 7.08 ppt, respectively. Conversely, at lower salinities of 1.79 ppt and 3.56 ppt, all fry survived beyond 12.0 d (Tebo and McCoy, 1964). Another investigation revealed that the 96-hour median tolerance limit of small LMB increased from 10.9 ppt to 13.4 ppt as body size ranged from 12-16 mm to 34-42 mm (Tebo and McCoy, 1964). This suggests a developmental variation in salinity tolerance. Similar patterns have been observed in other fish species. In the early life stages of yellow perch (Perca flavescens), yolk-sack larvae exhibited greater salinity tolerance at 12.9 ppt compared to the salinity tolerance of eggs at 12.1 ppt. In species like bigmouth buffalo (*Ictiobus cyprinellus*), black buffalo (*Ictiobus niger*), and channel catfish (*Ictalurus punctatus*), eggs demonstrated higher tolerance to saline waters than young juveniles. However, the salinity tolerance decreased at the time of hatching (Peterson and Meador, 1994). This research may contribute to understanding the changes in osmoregulatory capacity throughout the entire life history of LMB.

While traditionally recognized as a freshwater species, LMB has demonstrated adaptability to mesohaline environments. The distribution of fish in estuaries is believed to be primarily influenced by fluctuating salinity levels (Peterson and Ross, 1991). Fish typically adapt through the use of extracellular osmolytes at higher salinities, but they engage in a certain degree of hyperosmoregulation when salinity falls below 10 ppt (Madsen et al., 2015). Considering the potential impacts of freshwater salinization on LMB, understanding these dynamics may enhance the accuracy of predicting distribution and contribute to advancements in aquaculture technology.

## Conclusion

In summary, our data reveal that LMB eggs have an average diameter of 1.24 ± 0.09 mm. Both freshwater-activated mature eggs and naturally fertilized oocytes exhibit comparable osmotic homeostasis with similar diameters. Notably, the highest water excretion rate is observed at a salinity of 15.0 ppt when these eggs are cultured in saline water ranging from 0 ppt to 35.0 ppt. Under semi-natural conditions, the hatching rate of LMB embryos increases from the 1.0 ppt group to the 6.0 ppt group but significantly declines in the 8.0 ppt group. Yolk-sac LMB larvae exhibit survival for over 12 d in brackish water without feeding. However, the survival time gradually decreases as salinity increases from 0 ppt to 35.0 ppt. Logistic regression models effectively capture the relationship between lethal time and water salinity. Our study indicates that the osmoregulatory capability of LMB eggs may be primarily determined by the egg membrane, formed before ovulation. The observed difference in hyperosmoregulatory capacity between pre- and post-fertilization stages provides insights into the osmoregulatory mechanism of fish eggs. Moreover, at different ontogenetic stages, yolk-sac LMB larvae may adjust their hyperosmotic adaptability to acute salinity stress. Given the limited understanding of acute-phase regulation mechanisms in fishes, further investigations based on these findings have the potential to elucidate the process of hyperosmolality tolerance in a brackish environment.

## Acknowledgments

We would like to thank Mr. Zekun Chen for supplying with some largemouth bass samples and assisting with preparing the semi-field experiments in Foshan.

## Author contributions

DL and DY designed this research. DL performed the experiments and prepared the manuscript. All authors contributed to the article and approved the submitted version.

## Data availability statement

All data generated or analyzed during this study are included in this published article. The raw data supporting the conclusions of this article are available upon request.

## Declarations

## Ethical approval

All the animal study protocols were approved by the Laboratory Animal Ethics Committee of Pearl River Fisheries Research Institute, CAFS (LAEC-PRFRI-20160323).

## References

Adar RM, Safran SA (2020) Active volume regulation in adhered cells. Proc Natl Acad Sci USA 117: 5604–5609. 10.1073/pnas.1918203117

Andrea JR, Andrew KC, Irena FC, Erika JE, Peter AG, Pieter TJJ, et al. (2019) Emerging threats and persistent conservation challenges for freshwater biodiversity. Biol Rev 94: 849–873. 10.1111/brv.12480

Ankley GT, Bennett RS, Erickson RJ, Hoff DJ, Hornung MW, Johnson RD, et al. (2010) Adverse outcome pathways: A conceptual framework to support ecotoxicology research and risk assessment. Environ. Toxicol Chem 29: 730–741. 10.1002/etc.34

Baldisserotto B, (2019) Fish osmoregulation. CRC Press, Boca Raton

Ban M, Itou H, Nakashima A, Sada I, Toda S, Kagaya M, et al. (2022) The effects of temperature and salinity of hatchery water on early development of chum salmon (*Oncorhynchus keta*). Aquaculture 549: 737738. 10.1016/j.aquaculture.2021.737738

Chara O, Espelt MV, Krumschnabel G, Schwarzbaum PJ (2011) Regulatory volume decrease and p receptor signaling in fish cells: mechanisms, physiology, and modeling approaches. J Exp Zool Part A-Ecol Integr Physiol 315A: 175–202. 10.1002/jez.662

Coward K, Bromage NR, Hibbitt O, Parrington J (2002) Gamete physiology, fertilization and egg activation in teleost fish. Rev Fish Biol Fish 12: 33–58. 10.1023/A:1022613404123

Cui W, Ma A, Huang Z, Wang X, Liu Z, Xia D, Yang S, Zhao T (2020) Comparative transcriptomic analysis reveals mechanisms of divergence in osmotic regulation of the turbot scophthalmus maximus. Fish Physiol Biochem 46: 1519–1536. 10.1007/s10695-020-00808-6

Cunillera-Montcusí D, Beklioğlu M, Cañedo-Argüelles M, Jeppesen E, Ptacnik R, Amorim CA, et al. (2022) Freshwater salinisation: A research agenda for a saltier world. Trends Ecol Evol 37: 440–453. 10.1016/j.tree.2021.12.005

de Mutsert K, Cowan JH (2012) A before–after–control–impact analysis of the effects of a Mississippi river freshwater diversion on Estuarine Nekton in Louisiana, USA. Estuar Coasts 35: 1237–1248. 10.1007/s12237-012-9522-y

Devries D, Wright R, Glover D, Farmer T, Lowe M, Norris A, et al. (2015) Largemouth bass in coastal estuaries: A comprehensive study from the Mobile-Tensaw River Delta, Alabama. Am Fish Soc Symp 82: 297–309

Dumont JN, Brummet AR (1980) The vitelline envelope, chorion, and micropyle of *Fundulus heteroclitus* eggs. Gamete Res 3: 25–44. 10.1002/mrd.1120030105

Evans TG, Kültz D (2020) The cellular stress response in fish exposed to salinity fluctuations. J Exp Zool Part A-Ecol Integr Physiol 333: 421–435. 10.1002/jez.2350

Fridman S (2020) Ontogeny of the osmoregulatory capacity of teleosts and the role of ionocytes. Front Mar Sci 7: 709. 10.3389/fmars.2020.00709

Heidinger RC (1976) Synopsis of biological data on the largemouth bass, Micropterus salmoides (Lacepède) 1802. Food and Agriculture Organization of the United Nations, Rome

Herrera F, Bondarenko O, Boryshpolets S (2021) Osmoregulation in fish sperm. Fish Physiol Biochem 47: 785–795. 10.1007/s10695-021-00958-1

Hussein GHG, Chen M, Qi P, Cui Q, Yu Y, Hu W, et al. (2020) Aquaculture industry development, annual price analysis and out-of-season spawning in largemouth bass *Micropterus salmoides*. Aquaculture 519: 734901. 10.1016/j.Aquaculture.2019.734901

Katelyn ML, Jeffrey EH (2021) Predicting successful reproduction and establishment of non-native freshwater fish in peninsular Florida using life history traits. J Vertebr Biol 70: 21041. 10.25225/jvb.21041

Keup L, Bayless J (1964) Fish distribution at varying salinities in Neuse River Basin, North Carolina Chesapeake Sci 5: 119–123. 10.2307/1351370

Kültz D (2012a) 2 - osmosensing, in: McCormick, SD, Farrell, AP, Brauner, CJ (Eds.), Fish Physiology-Euryhaline Fishes. Academic Press, Waltham, pp 45–68

Kültz D (2012b) The combinatorial nature of osmosensing in fishes. Physiology 27: 259–275. 10.1152/physiol.00014.2012

Kültz D (2015) Physiological mechanisms used by fish to cope with salinity stress. J Exp Biol 218: 1907–1914. 10.1242/jeb.118695

Kumar V, Sheoran OP, Rani S, Malik K (2020) Development of a web-based tool for probit analysis to compute LC50/LD50/GR50 for its use in toxicology studies. J Appl Nat Sci 12: 641–646. 10.31018/jans.v12i4.2408

Lionetto MG, Giordano ME, De Nuccio F, Nicolardi G, Hoffmann EK, Schettino T (2005) Hypotonicity induced k^+^ and anion conductive pathways activation in eel intestinal epithelium. J Exp Biol 208: 749–760. 10.1242/jeb.01440

Lowe MR, DeVries DR, Wright RA, Ludsin SA, Fryer BJ (2009) Coastal largemouth bass (*Micropterus salmoides*) movement in response to changing salinity. Can J Fish Aquat Sci 66: 2174–2188. 10.1139/F09-152

Lu G, Yao Z, Lai Q, Gao P, Zhou K, Zhu H, et al. (2022) Growth performance, blood parameters and texture characteristics of juvenile largemouth bass (*Micropterus salmoides*) exposed to highly saline-alkaline water. Progress in Fishery Sciences 43: 1–11. 10.19663/j.issn20959869.20220112002

Madsen SS, Engelund MB, Cutler CP (2015) Water transport and functional dynamics of aquaporins in osmoregulatory organs of fishes. Biol Bull 229: 70–92. 10.1086/BBLv229n1p70

Marshall WS, Howard JA, Cozzi RRF, Lynch EM (2002) Nacl and fluid secretion by the intestine of the teleost fundulus heteroclitus: involvement of cftr. J Exp Biol 205: 745–758. 10.1242/jeb.205.6.745

Matthews MD, Sakmar JC, Trippel N (2012) Evaluation of hydrogen peroxide and temperature to control mortality caused by saprolegniasis and to increase hatching success of largemouth bass eggs. N Am J Aquacult 74: 463–467. 10.1080/15222055.2012.676608

McCormick SD, Farrell AP, Brauner CJ (2013) Fish physiology: Euryhaline fishes. Academic Press, Waltham

Meador MR, Kelso, WE (1989) Behavior and movements of largemouth bass in response to salinity. Trans Am Fish Soc 118: 409–415. 10.1577/15488659(1989)118<0409:BAMOLB>2.3.CO;2

Meador MR, Kelso, WE (1990) Growth of largemouth bass in low-salinity environments. Trans Am Fish Soc 119: 545–552. 10.1577/1548-8659(1990)119<0545:GOLBIL>2.3.CO;2

Meador MR, Kelso, WE (1990) Physiological responses of largemouth bass, *Micropterus salmoides*, exposed to salinity. Can J Fish Aquat Sci 47: 2358–2363. 10.1139/f90-262

Miller KD (1971). Spawning and early life history of largemouth bass (Micropterus salmoides) in Wahweap Bay, Lake Powell. Dissertation, Utah State University

Norris AJ, DeVries DR, Wright RA, (2010) Coastal estuaries as habitat for a freshwater fish species: Exploring population-level effects of salinity on largemouth bass. Trans Am Fish Soc 139: 610–625. 10.1577/T09-135.1

Peterson MS (1988) Comparative physiological ecology of centrarchids in hyposaline environments. Can J Fish Aquat Sci 45: 827–833. 10.1139/f88-100

Peterson MS, Meador MR (1994) Effects of salinity on freshwater fishes in coastal plain drainages in the southeastern U.S. Rev Fish Sci 2: 95–121. 10.1080/10641269409388554

Peterson MS, Ross ST (1991) Dynamics of littoral fishes and decapods along a coastal river-estuarine gradient. Estuar Coast Shelf Sci 33: 467–483. 10.1016/0272-7714(91)90085-P

R Core Team (2022) R: A language and environment for statistical computing. R Foundation for Statistical Computing, Vienna, Austria. https://www.R-project.org/. Accessed 11 June 2022

Rizzo E, Bazzoli N (2020) Chapter 13 - Reproduction and embryogenesis. In Baldisserotto B, Urbinati EC, Cyrino JEP (ed) Biology and physiology of freshwater neotropical fish. Academic Press, London, pp 287-313

Rueden CT, Schindelin J, Hiner MC, DeZonia BE, Walter AE, Arena ET, et al. (2017) ImageJ2: ImageJ for the next generation of scientific image data. BMC Bioinf 18: 529. 10.1186/ s12859-017-1934-z

Schoots AFM, Sackers RJ, Overkamp PSG, Denucé, JM (1983) Hatching in the teleost *Oryzias latipes*: Limited proteolysis causes egg envelope swelling. J Exp Zool 226: 93–100. 10.1002/jez.1402260112

Segret E, Cardona E, Skiba-Cassy S, Cachelou F, Bobe J (2022) Effect of a low water concentration in chloride, sodium and potassium on oocyte maturation, oocyte hydration, ovulation and egg quality in rainbow trout. Aquaculture 546: 737374. 10.1016/j.aquaculture.2021.737374

Sørensen SR, Butts IAE, Munk P, Tomkiewicz J (2016) Effects of salinity and sea salt type on egg activation, fertilization, buoyancy and early embryology of European eel, *Anguilla anguilla*. Zygote 24: 121–138. 10.1017/S0967199414000811

Takei Y, Hwang P (2016) 6 - homeostatic responses to osmotic stress. In: Schreck CB, Tort L, Farrell AP, Brauner CJ (ed) Fish Physiology. Academic Press, Waltham, pp 207–249

Tebo LB, McCoy EG (1964) Effect of sea-water concentration on the reproduction and survival of largemouth bass and bluegills. Prog Fish Cult 26: 99–106. 10.1577/15488640(1964)26[99:EOSCOT]2.0.CO;2

Tidwell JH, Coyle SD, Bright LA (2019) Largemouth bass aquaculture. 5m Publishing, Sheffield

Wang X, Kültz D (2017) Osmolality/salinity-responsive enhancers (osres) control induction of osmoprotective genes in euryhaline fish. Proc Natl Acad Sci USA 114: E2729–E2738. 10.1073/pnas.1614712114

Wang Y, Chen F, He J, Xue G, Chen J, Xie P (2021) Cellular and molecular modification of egg envelope hardening in fertilization. Biochimie 181: 134–144. 10.1016/j.biochi.2020.12.007

Yi H, Chen X, Liu S, Han L, Liang J, Su Y et al. (2021) Growth, osmoregulatory and hypothalamic– pituitary–somatotropic (HPS) axis response of the juvenile largemouth bass (*Micropterus salmoides*), reared under different salinities. Aquacult Rep 20: 100727. 10.1016/ j.aqrep.2021.100727

York-Andersen AH, Wood BW, Wilby EL, Berry AS, Weil TT (2021) Osmolarity-regulated swelling initiates egg activation in *Drosophila*. Open Biol 11: 210067. 10.1098/rsob.210067

Zelazowska M, Halajian A (2019) Previtellogenic oocytes of South African largemouth bass *Micropterus salmoides* Lacepede 1802 (*Actinopterygii, Perciformes*) - the Balbiani body, cortical alveoli and developing eggshell. J Morphol 280: 360–369. 10.1002/jmor.20948

